# A finished, telomere-to-telomere assembly for *Botrytis cinerea* strain B05.10

**DOI:** 10.64898/2026.10.20.760327

**Authors:** Jan A.L. van Kan, Dian Zheng, Frank P.J. Pieterse, Xiaoqian Shi-Kunne

## Abstract

We present an improved genome assembly of Botrytis cinerea strain B05.10 which serves as a reference isolate for most labs active in the Botrytis research. The existing assembly dating from 2017 was extended by adding 8 telomeric regions and a small number of minor errors was corrected. The new assembly contains telomeres on each chromosome end, except for Chr4 carrying the rDNA repeat. The unannotated assembly is deposited in NCBI and manual curation of gene models is ongoing. The new assembly will be hosted on Ensembl-Fungi after completion of the structural curation.

## Introduction

The fungus *Botrytis cinerea* is an important plant pathogen for economic and scientific reasons and is the most extensively investigated necrotrophic phytopathogenic fungus (Dean et al., 2012). Besides studies on its infection mechanisms (Bi et al., 2022) and the development of disease control methods through breeding (You et al., 2025), cultural practices (Mundy et al., 2022) or biocontrol (Bourgeay and Bardin, 2026), the responses of the fungus to light and circadian rhythm, as well as its sexual reproduction have been studied (Schumacher, 2017; Hevia et al., 2015; Rojas et al., 2023; Rodenburg et al., 2018).

The fungus can infect over 1500 host plant species from >450 families and it thereby has one of the largest host ranges recorded for any fungal plant pathogen. Despite its broad host range, there is evidence for the occurrence of host preference and host adaptation in the *B. cinerea* population, with isolates sampled from grape or tomato being more virulent on the host plant from which they were sampled (Mercier et al., 2019; Mercier et al., 2021). Host preference was shown to be associated with the tolerance of a host-adapted isolate to antifungal compounds produced by the host plant (You et al., 2024).

Strain B05.10 has served as the reference for *B. cinerea* research since the late 1990’s and is the most commonly used strain in many dozens of labs across the globe. The origin of strain B05.10 has been unclear and somewhat mysterious for many people. It was obtained in experiments by Büttner et al. (1994) who exposed isolate SAS56 to sublethal doses of benomyl, because there were indications that isolate SAS56 could be polyploid or aneuploid. The benomyl treatment was applied to distort chromosomal segregation during mitosis with the aim to cause the outgrowth of sectors with altered ploidy levels and thereby yield stable haploid derivatives for genetic analyses. Emerging hyphal sectors were propagated and their nuclear DNA content was quantified (Büttner et al., 1994). One of the strains obtained from these experiments was B05.10, which had an apparent lower DNA content per nucleus (as compared to SAS56), was very stable in culture, virulent on multiple hosts and well transformable. It was only 2-3 years later that it turned out that B05.10 was not derived from SAS56, as there were very significant sequence polymorphisms between both strains, as well as several transposon insertions that were present in one but not the other strain. The conclusion was that B05.10 could not be a derivative of SAS56 and therefore must have been a benomyl-resistant contaminant in the experiment. Indeed a single point mutation in the β-tubulin gene that confers resistance to benomyl (van der Vlugt-Bergmans et al., 1993) was present in B05.10 but not in SAS566. The exact source of the contamination has never been revealed.

Genome sequences have become an essential resource in fungal research in the past decades. The first effort to sequence the genome of B05.10 was carried out by Syngenta at the Torrey Mesa Research Institute in La Jolla (California) in the late 1990’s. Data were generated by Sanger sequencing of a plasmid library and sequence data were initially not public. Around the turn of the century, Syngenta agreed to share information by performing Blast searches with queries that researchers from the *Botrytis* community would submit. The Blast output was shared and gave the researchers a first glimpse of the exact sequence and genomic environment of their gene of interest. Access to the genome information was interrupted when Syngenta terminated its target gene discovery research in *Botrytis*. In 2005, Syngenta agreed to transfer the raw sequence data from the 1990’s to the Broad Institute in Cambridge (Massachussets). A group headed by Christina Cuomo assembled the Sanger reads into the genome that was eventually published, along with genomes of *B. cinerea* strain T4 and of *Sclerotinia sclerotiorum* (Amselem et al. 2011). This publication contained extensive descriptions of various properties and biological features of both fungal species that were better understood from their genomic information. However, the fragmentation of the genome into 588 scaffolds and the low (4.5X) coverage prevented a detailed insight into chromosome architecture of the strain.

An updated version of the *B. cinerea* B05.10 and T4 genomes based on Illumina reads was communicated in 2012 (Staats and van Kan, 2012) which served as a stepping stone to the description of a gapless genome of B05.10, based on Illumina and PacBio data, and supported by an optical and genetic map (van Kan et al., 2017). This gapless genome, however, was still incomplete because it lacked eight telomeric regions that were present in the data, but could not accurately be assigned to chromosomes. From the optical map information, it was inferred that a total of 847 kbp was missing from the assembly, including 580 kbp of rDNA repeats at the start of Chr4. Here we report the use of Nanopore technology to obtain a finished telomere-to-telomere assembly for B05.10, with the exception of the chromosome arm carrying the rDNA repeat.

## Methods

### DNA isolation and sequencing

Mycelium was collected from a culture on Malt Extract agar overlaid with cellophane, and freeze-dried. The mycelium was grinded in a Retsch machine with two glass beads. A part of the powder was extracted with Puregene cell lysis solution (Qiagen) supplemented with Proteinase K and RNase A according to manufacturer’s prescription. To the lysate we added Puregene protein precipitation solution (Qiagen), incubated for 15 minutes on ice and centrifuged the mixture at 17000g for 2 min at room temperature. Supernatant was transferred to a new tube and centrifuged again at 20000g for 10 min in the cold room. The resulting supernatant was collected and one volume of Qiagen QBT buffer was added. The sample was thoroughly mixed by repeated inversion of the tube and applied to a Qiagen Genomic Tip 20, pre-equilibrated with QBT. The column was washed with 3 volumes of buffer QC and subsequently eluted with 2 × 1ml of buffer QF. The eluted DNA was equally divided over two 2 ml tubes, and precipitated by adding 0.7 ml isopropanol followed by centrifugation for 10 min at 20000g in the cold room. The pellet was washed twice with 70% EtOH, air-dried for ~10 min and redissolved in TE buffer overnight in the cold room. The DNA was quantified by Qubit and run on an agarose gel for size estimation.

Nanopore library construction and sequencing was performed at IGA Technologies in Udine (Italy). DNA was size-fractionated on a Blue Pippin cassette (cut-off = 15 kbp) prior to library construction, the library was sequenced on a Nanopore Promethion machine.

### Improvement of the B05.10 genome assembly

The publicly available *B. cinerea* B05.10 genome assembly (van Kan et al., 2017) was used as starting reference for manual improvement of the chromosome ends. An independent assembly of B05.10 was generated from ONT reads using NECAT (version v0.0.1_update20200803; Chen et al., 2021). The NECAT assembly was subsequently used as a source of candidate sequences for correcting and extending chromosome ends in the publicly available assembly. Both the reference assembly and the NECAT assembly were evaluated against the raw ONT reads to assess the completeness and accuracy of the assembled chromosome ends.

For both assemblies, the B05.10 ONT reads were mapped using minimap2 (version 2.30-r1287) with the “map-ont” preset, with secondary alignment disabled (Li et al., 2018). A custom filtering script was subsequently applied to retain high-confidence ONT read alignment. Unmapped, secondary, and supplementary alignments were removed. Reads were retained only when they had a mapping quality (MAPQ) of at least 20, at least 90% of the original read length aligned to the assembly, and no more than 5% of the read length was soft- or hard-clipped. The filter alignments were subsequently sorted and indexed using SAMtools (version 1.23; Danecek et al., 2021) for visualization in IGV (Robinson et al., 2011; Thorvaldsdóttir et al., 2013).

The reference assembly was first inspected to identify chromosomes lacking complete telomeric sequences. For each such chromosome, the terminal 20-50 kb was examined in IGV to determine whether ONT reads extended beyond the assembled chromosome end. The corresponding chromosome ends in the NECAT assembly were then examined using the ONT read alignments to determine whether additional sequence was present and supported by the raw sequencing data. Candidate extensions identified in the NECAT assembly were incorporated into the reference assembly only when they were supported by multiple independent ONT reads and showed consistent read alignment. The ONT reads alignment to the NECAT assembly were used to verify that candidate sequence was supported by the sequencing data and was not an artifact of the NECAT assembly.

The resulting modified B05.10 assembly was subsequently assessed manually in IGV and by Tapestry (version 1.0.1; Davey et al., 2020) for the presence of telomeric sequences.

Dotplots were generated using D-Genies (Cabanettes and Klopp, 2018).

## RESULTS

### DNA isolation for Nanopore sequencing

It is imperative to use high molecular weight (HMW) DNA of excellent quality for preparing Nanopore libraries. The isolation of HMW DNA from *B. cinerea* is notoriously challenging. In liquid cultures the fungus produces a slimy α1-3,1-6 glucan polymer, called cinerean (Stahmann et al., 1995) that co-purifies with the DNA to significant extent. Therefore liquid cultures are undesirable for this purpose. Better DNA quality was obtained when the fungus was grown on agar medium on a cellophane sheet, from which mycelium could be scraped. Nevertheless, there still was cinerean at the interphase of the hyphae with cellophane; it was beneficial to submerge the mycelial mat upside down in water and gently rub the slime off. The mycelium was subsequently freeze-dried and grinded into powder in a Retsch shaker. The powder should remain relatively coarse, grinding for a very long period resulted in fine powder that yielded DNA of lower molecular weight. The lysis and DNA isolation was done with a classical CTAB method and the resulting DNA was purified on a Qiagen genomic tip, which works on gravity-based elution. Purification on various types of spin columns resulted in DNA with significantly lower molecular weight, even when the column was supposed to be suitable for HMW DNA. Running the sample on an agarose gel showed that the majority of the DNA was >25 kbp in size. Storage of the DNA in unfrozen state helped to retain the proper length. Blue Pippin size separation was performed (cutoff=15 kbp) prior to construction of the Nanopore sequencing library.

### Sequencing and assembly

The sequencing yielded 315,306 reads (5.79 Gbp in total) with a median read length of 15,908 bp and median read quality of 20.0. The longest read was 170,198 bp in length. The overall read coverage for the entire B05.10 genome (~43 Mbp) was therefore 135 X.

Mapping the long (>30kbp) reads to the existing gapless reference genome (van Kan et al., 2017) revealed assembly errors for the 5’-telomeric region of Chr2, which appeared to be composed of a mixture of multiple haplotypes of similar, non-identical sequences, some of which carried additional repetitive elements. In order to join the correct telomeric sequence to Chr2, we manually selected the 5 longest mapping Nanopore reads (>50 kbp) that contained distal sequences that were definitely unique to Chr2. The consensus of these reads was used to replace the previous 5’-telomeric region of Chr2, resulting in an increase in length of this region by ~7kbp.

For the 8 telomeric regions missing in the reference genome (van Kan et al., 2017), we mapped the long reads to the existing gapless reference and manually selected reads that contained extensions towards the telomere. The clipped (non-mapping) regions of such reads were used to generate a consensus and inserted in front or downstream of the existing reference sequence. Table 1 summarizes the adjustments that were made to the previous assembly.

**Table 1.** Sequence adjustments in new assembly.

|  | Missing sequence<sup>a</sup> | Alterations | Comment |
| --- | --- | --- | --- |
| Chr2 | 0 | 6,904 bp added | Deconvolution multiple haplotypes, Homology to telomere Chr10R |
| Chr5 | 50 kbp (5') | 60.5 kbp added | Homology to telomere Chr16R |
| Chr5 | 22 kbp (3') | 26.5 kbp added | Homology to telomere Chr7F |
| Chr7 | 28 kbp (5') | 34.4 kbp added | Homology to telomere Chr5R |
| Chr8 | 0 | 13.3 kbp added | Insertion in 5'-telomeric region |
| Chr9 | 25 kbp (5') | 29.5 kbp added | Homology to telomere Chr10R |
| Chr10 | 42 kbp (3') | 28.5 kbp added | Homology to telomeres Chr2F, 9F, 11F, 14F |
| Chr11 | 25 kbp (5') | 27.6 kbp added | Homology to telomere Chr10R |
| Chr14 | 32 kbp (5') | 33.1 kbp added | Homology to telomere Chr10R |
| Chr16 | 43 kbp (3') | 50 kbp added | Homology to telomere Chr5F |

<sup>a</sup> Estimate of missing sequence based on optical map (van Kan et al., 2017)

### Final quality check of the assembly

The resulting assembly (Figure 1) was inspected by mapping the long reads with high stringency and inspecting the IGV plot for errors and inconsistencies. Read coverage was generally >100 across the chromosomes, with somewhat lower coverage towards the telomeric regions, especially in case of newly added telomeric fragments which were highly similar to telomeres of other chromosomes. No conflicts were detected that would justify further polishing or other adjustments.

**Figure 1.**
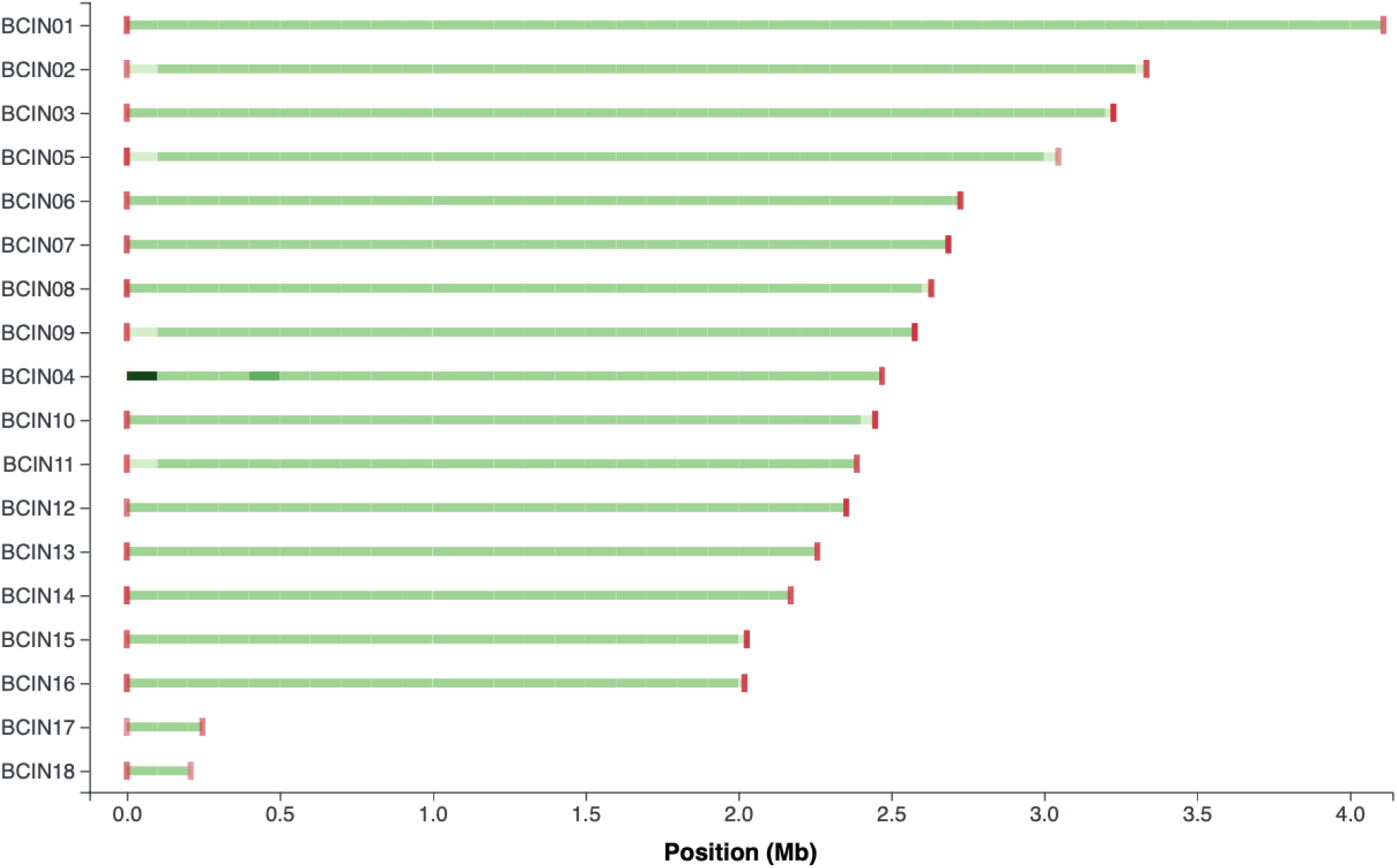
Tapestry visualization of genome assembly contigs with read mapping density. The figure shows a representation of the contigs of the genome assembly, with the green color density indicating the number of reads that have been mapped to each contig. The contigs are ordered by length, with the longest contigs at the top. The red bars indicate telomeric repeats. Note that chromosome BIN04 is in reality ~ 550kb longer because it contains ~65 copies of the rDNA repeat.

### Duplications of telomeric regions

After properly adding the telomeric regions to their corresponding chromosomes, we examined the cause for the earlier problems in the assembly by van Kan et al. (2017). A dotplot was generated for a concatenated sequence that comprised the 50 kbp telomeric regions (5’and 3’) of each of the 18 chromosomes (Figure 2). The dotplot shows that the 5’-telomeric region of Chr2 is homologous to the 3’-telomeric region of Chr10; the 5’-telomeric region of Chr5 is homologous to the 3’-telomeric region of Chr16, and the 3’-telomeric region of Chr5 is homologous to the 5’-telomeric region of Chr7. Notably, the most widespread homology was observed between the 3’-telomeric region of Chr10 and the 5’-telomeric regions of four distinct chromosomes, i.e. Chr2, Chr9, Chr11 and Chr14. All homologies were in inverted orientation, implying that there are several shared (sub)families of telomeric region sequences. The homology did not extend (much) beyond the 50 kbp boundary, suggesting that 50kbp is a reasonable estimate for the maximal length of true telomeric regions in this fungus.

**Figure 2.**
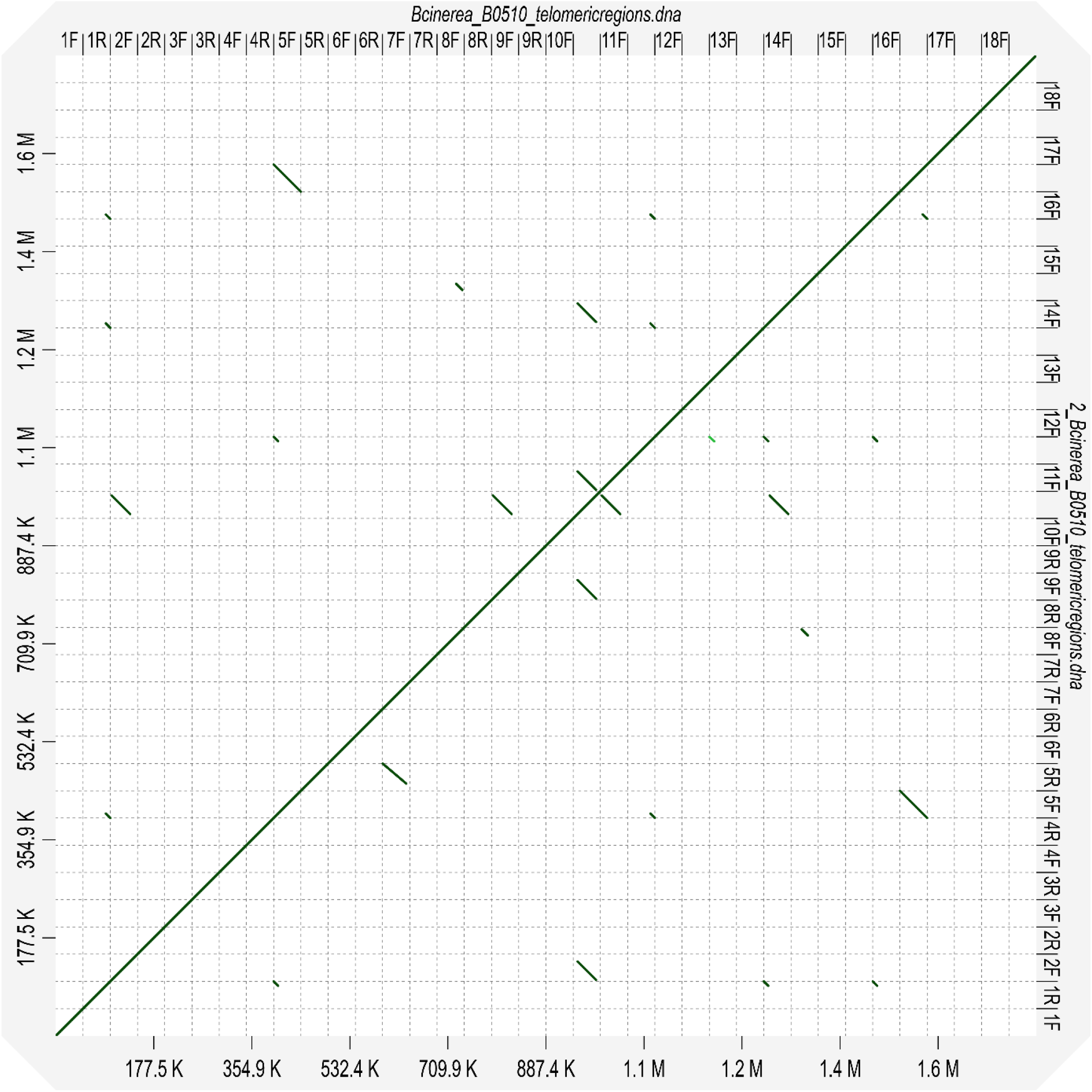
Dotplot of a concatenated sequence comprising 50 kbp stretches of 5’- and 3’-telomeric regions of the 18 *B. cinerea* chromosomes, plotted against itself. Chromosomes are numbered 1-18, F refers to the 5’-telomeric region while R refers to the 3’-telomeric region.

### Gene prediction and annotation

The gene models predicted on the earlier reference assembly (van Kan et al., 2017) were manually curated by multiple members of the *Botrytis* community. This curation made use of RNAseq data that were available at the time, which represented a wide range of developmental stages and treatments. Efforts were made to define the transcription start and stop sites and identify splice variants (also in UTR’s). However, the quality of RNAseq data in 2016 is nowadays considered inferior, as the reads were shorter and not strand-specific. Old gene models have been transferred to the new assembly and will undergo a new round of curation by a single person, using more recent strand-specific RNAseq datasets. Once manual curation is completed, the new gene models will be deposited in NCBI and the annotated genome will be hosted on Ensembl-Fungi.

## DISCUSSION

The earlier gapless reference genome (van Kan et al., 2017) was of high quality because of its validation by an optical and genetic map, however it lacked eight telomeric regions due to the significant homology between telomeric regions of distinct chromosomes. Several strategies were used to add the missing telomeric regions and check the structural accuracy of the telomeric region assembly, as well as polish minor base call errors. This procedure required quite some manual intervention because of the significant duplications between telomeric regions, which were up to 45 kb in length (between Chr 5 and 16). The current assembly reflects a complete and correct telomere-to-telomere (T2T) genome sequence for B05.10. The rDNA region that is present at the 5’-end of Chr4 could not be assembled as it contains ~65 copies of the rDNA unit, based on the estimate from the optical map (van Kan et al., 2017).

It will be of interest to analyse telomere dynamics in other *B. cinerea* strains on a pangenomic scale. It was reported by Valero-Jiménez et al. (2020) that the synteny between different *Botrytis* species was remarkably high, even between species that are members of distinct phylogenetic clades within the genus, which separated ~5 MYA. For example, *B. aclada* contains 11 core chromosomes (of the 16) which are highly syntenic to their counterparts in *B. cinerea* B05.10 over >95% of their length, and the few rearrangements between the 11 core chromosomes of these two species predominantly comprise six sub-telomeric rearrangements (Valero-Jiménez et al., 2020).

The currently ongoing systematic curation of gene models will yield even more accurate insights into transcript structures, including transcription start and polyadenylation sites, as well as differential splicing events in different fungal cell/tissue types, or under distinct conditions. The new assembly will also lay the foundation for annotating lncRNAs. Eventually, all transcript data should be integrated with information about chromatin architecture and dynamics, which are currently understudied in *B. cinerea*. All these resources will contribute to maintaining *B. cinerea* as the best studied necrotrophic plant pathogenic fungus.

## Data availability

This Genome has been deposited at DDBJ/ENA/GenBank under the accession AAID00000000. The version described in this paper is version AAID03000000.

The raw read data used for this assembly are part of BioProject accession number PRJNA15632, the reads are deposited under accession SRR40741277.

## Acknowledgments

The authors acknowledge partial financial support by the 4TU-FairData programme awarded to DZ. The research of DZ is sponsored by the China Scholarship Council. The research of FPJP is supported by NWO grant OCENW.M.22.213.

